# Real-space quantum-based refinement for cryo-EM: Q|R#3

**DOI:** 10.1101/2020.05.25.115386

**Authors:** Lum Wang, Holger Kruse, Oleg V. Sobolev, Nigel W. Moriarty, Mark P. Waller, Pavel V. Afonine, Malgorzata Biczysko

## Abstract

Electron cryo-microscopy (cryo-EM) is fast becoming a major competitor to X-ray crystallography especially for large structures that are difficult or impossible to crystallize. While recent spectacular technology improvements are leading to significantly higher resolution of three-dimensional reconstructions, the average quality of cryo-EM maps is still on the low-resolution end of the range compared to crystallography. A long-standing challenge for atomic model refinement has been the production of stereochemically meaningful models for this resolution regime. Here we demonstrate how including accurate model geometry restraints derived from *ab initio* quantum-chemical calculations (HF-D3/6-31G) can improve the refinements of an example structure (chain A of 3j63). The robustness of the procedure is tested for additional structures with up to 7k atoms (3a5x, and chain C of 5fn5) by means of the less expensive semi-empirical (GFN1-xTB) model. Necessary algorithms enabling real-space quantum refinement are implemented in the latest version of *qr.refine* and are described herein.

**Synopsis:** The implementation of quantum-based real-space refinement in *qr.refine* is described.

## 1. Introduction

Improvements in cryo-EM technology resulted in a rapidly increasing number of three-dimensional reconstructions of 4.5 Å and better (Kühlbrandt, 2014, Henderson, 2015, Nogales, 2015, Orlov *et al.*, 2017, Baldwin *et al.*, 2018) allowing interpretation of corresponding maps in terms of atomic models. This prompted the active development of map improvement (Terwilliger, Sobolev, *et al.*, 2018; Terwilliger, Ludtke, *et al.*, 2019), model building (Terwilliger, Adams, *et al.*, 2018, 2019), refinement (Afonine, Poon, *et al.*, 2018) and validation methods (Afonine, Klaholz, *et al.*, 2018). It is a common knowledge that atomic model refinement can be challenging at resolutions worse than about 3-3.5 Å (Headd *et al.*, 2012). This is because beyond this resolution the amount of detail in the experimental data is insufficient to resolve structural features of proteins such as secondary-structure or rotameric states of amino-acid residues at an atomic level. This in turn requires extra information to be used in refinement, for example: Ramachandran plot, rotamer, secondary-structure or reference model restraints (for a summary, see for example Afonine, Poon, *et al.,* 2018). Quantum-mechanical (QM) calculations generate model geometries *ab initio* and thus do not rely on empirical libraries, such as the commonly used Monomer Library (Vagin & Murshudov, 2004; Vagin *et al.*, 2004) nor any other ad hoc restraints. Historically, quantum refinements are associated with either impractical runtimes even for small proteins or methods that focus on QM calculations only for the macromolecular region of interest, such as a ligand-binding pocket or an enzyme’s active site, while treating the rest of the molecule with a classical approach (Ryde, 2003; Ryde & Nilsson, 2003b, a; Nilsson *et al.*, 2004; Borbulevych *et al.*, 2014). Recently we have demonstrated that the entire atomic model can benefit from a full QM treatment (Zheng, Reimers, *et al.,* 2017) being the first to enable full protein crystal structure refinement using QM based geometry gradients in place of the gradients from classical restraints (Zheng, Moriarty, et al., 2017).

Quantum Refinement (Q|R; www.qrefine.com) is an international collaborative project with the goal to create easy to use software for full structure refinement using gradients derived from quantum-based calculations. Q|R is built upon the CCTBX library (Grosse-Kunstleve *et al.*, 2002) and operates as a plugin to the *Phenix* software (Liebschner *et al.*, 2019). It uses Atomic Simulation Environment (ASE) adaptors (Bahn *et al.*, 2002) to interface to a number of available QM packages such as TeraChem (Ufimtsev *et al.*, 2009; Titov *et al.*, 2012), Orca (Neese, 2012), xtb (Grimme *et al.*, 2017; Bannwarth *et al.*, 2019), MOPAC (Stewart, 2016), TURBOMOLE (Furche *et al.*, 2014), Gaussian (Frisch *et al.*, 2016) and more. Q|R is a work in progress with set challenges that we address as the project progresses. Describing fundamentals (Zheng, Reimers, *et al.*, 2017), proving concept (Q|R#1, Zheng, Moriarty, *et al.*, 2017), handling crystallographic symmetry (Q|R#2, Zheng *et al.*, 2020), supporting cryo-EM (Q|R#3, this manuscript), alternative conformations (Q|R#4, manuscript in preparation) and benchmarking different QM methods to optimize speed and accuracy of calculations, are examples of such challenges and milestones.

Recent development of real-space methods and tools in CCTBX enabled the implementation of classic real-space refinement in *Phenix* (Afonine, Poon, *et al.*, 2018) specifically designed for cryo-EM but also supports crystallographic data. Since Q|R is built on CCTBX and is tightly integrated with *Phenix*, it was possible to re-use the key components of CCTBX to implement quantum-based real-space refinement in Q|R. Herein we describe the implementation details and demonstrate the utility of the method and reliability of geometry restraints derived from *ab initio* quantum-chemical calculations using select examples.

## 2. Methods

### 2.1. Real-space refinement in Q|R

Similarly to reciprocal-space refinement (Afonine *et al.*, 2012), the refinement target function is a composite of two terms

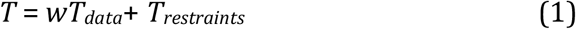

where *T_data_* scores the model-to-data fit, *T_restraints_* represents *a priori* information about the problem and *w* is an empirical weight factor that allows to maximize model-to-data fitting while enforcing the model to conform to the prior knowledge. Except for rare cases when ultra-high resolution (0.7 Å or better^1^) data are available (Dauter *et al.*, 1992; Bönisch *et al.*, 2005; Wang *et al.*, 2007; Hirano *et al.*, 2016), refinement of macromolecular models always requires using *T_restraints_* to compensate for the limited quality of experimental data (e.g. finite resolution).

In contrast to reciprocal-space refinement, the experimental data used in (1) are not structure factor amplitudes or intensities but a real-space map. This allows the calculations to be done locally, which in turn enables faster calculations and larger refinement convergence radii (Afonine, Poon, *et al.*, 2018). Indeed, in reciprocal space refinement all atoms and all *hkl* Miller indices corresponding to measured data are required in order to match model-calculated structure factors with measured ones. In real-space refinement the three-dimensional reconstruction, the map, represents the experimental data and atoms can move locally to optimize fit to the map. This, for example, allows use of only a fraction of the atomic model to find the optimal weight *w* in (1), which is computationally efficient compared to using the whole model as implemented in *phenix.real_space_refine* and described by Afonine, Poon, *et al.* (2018).

Q|R uses the *cctbx.iotbx* module of CCTBX to import an MRC formatted map (Winn *et al.*, 2011) and atomic model in PDB (Bernstein *et al.*, 1977) or mmCIF formats (Adams *et al*., 2019). The *cctbx.maptbx* module is used to perform calculation of (1) and its gradients as described in Afonine, Poon, *et al.* (2018). The *cctbx.maptbx.real_space_refinement_simple* module is used to carry out the optimization of (1). In this work, it is assumed that a cryo-EM map is sampled in a P1 symmetry box with the model situated in its center and, unlike for crystallography, no periodicity needs to be taken into account. Optimization of the weight *w* in (1) uses the same idea as in *phenix.real_space_refine* but is implemented differently owing to specifics of QM calculations. *phenix.real_space_refine* systematically probes the weight in the range from very small (least emphasis on the data) to very large (most emphasis on the data). This can cause problems in *qr.refine* because a trial refinement run with a very large weight is likely to yield a very geometrically distorted atomic model that in turn is likely to impede QM calculations. Therefore in quantum refinement the order of trial weights is important. Here the weights are sampled from smallest to largest and for each trial weight several parameters are monitored. These are, the root-mean-squared deviation (RMSD) of bond lengths from “ideal” library values (Vagin & Murshudov, 2004; Vagin *et al*., 2004) as in the *phenix.real_space_refine*, and additionally the percentage of residues in the favored region of the Ramachandran plot (% favored), percentage of rotamer outliers, Cβ deviation and the Clash-score. These define the overall quality of the model with deviation growing as the trial weight increases; the weight search terminates once the bond RMSD reaches the predefined threshold or any other parameter worsen. The refinement was performed with the *qr.refine* version v1.0-176 coupled to the Phenix 1.17.

### 2.2. Test model selection, preparation, refinement setup and analysis

Three atomic models and corresponding maps have been selected from the Protein Data Bank (wwPDB consortium, 2019) and Electron Microscopy Data Bank (Lawson et al., 2011) with a resolution range between 3.8 and 4.3 Å (Table 1). The 3a5x model was used as is, while only chains A and C were used from 3j63 and 5fn5, respectively. Each model was subject to a fully automated preparation for quantum refinement using the *qr.finalise* tool (Zheng, Moriarty, *et al.*, 2017) of the Q|R package; this includes completing missing side chains and adding all theoretically possible (but missing) hydrogen atoms. For partial models (3j63 and 5fn5) a box with the map around corresponding chains was extracted using the *phenix.map_box* tool (Liebschner *et al.*, 2019).

**Table 1.** Summary of data, model and model to data fit statistics for initial model (Initial) and models after refinement using CCTBX (with tight and loose geometry restraints) and quantum restraints derived from TeraChem (TC) or GFN1-xTB (xtb). CC_mask_ reports the correlation between the model-generated and experimental maps. Atom/residue counts for 5nf5 and 3a5x are shown for partial models actually used in refinements.

| PDB/EMDB |  | 3j63/5830 |  |  |  |  | 5fn5/3240 |  |  |  | 3a5x/1641 |  |  |  |
| --- | --- | --- | --- | --- | --- | --- | --- | --- | --- | --- | --- | --- | --- | --- |
| Atoms/residues |  | 1436/91 |  |  |  |  | 3787/243 |  |  |  | 7194/494 |  |  |  |
| Resolution (Å) |  | 3.8 |  |  |  |  | 4.3 |  |  |  | 4 |  |  |  |
| Metric | Model | Initial | CCTBX<br>(loose) | CCTBX<br>(tight) | xtb | TC | Initial | CCTBX<br>(loose) | CCTBX<br>(tight) | xtb | Initial | CCTBX<br>(loose) | CCTBX<br>(tight) | xtb |
| CC<sub>mask</sub> |  | 0.67 | 0.74 | 0.67 | 0.61 | 0.67 | 0.64 | 0.72 | 0.66 | 0.64 | 0.31 | 0.55 | 0.32 | 0.33 |
| RMSD | Bonds (Å) | 0.004 | 0.015 | 0.002 | 0.012 | 0.015 | 0.012 | 0.015 | 0.002 | 0.009 | 0.014 | 0.015 | 0.001 | 0.010 |
|  | Angles (°) | 1.22 | 1.30 | 0.48 | 1.82 | 1.61 | 2.00 | 1.40 | 0.50 | 1.57 | 1.65 | 1.62 | 0.39 | 1.82 |
| Rama.<br>plot<br>(%) | Favored | 85.4 | 55.1 | 96.6 | 98.9 | 100 | 90.9 | 61.0 | 89.2 | 94.6 | 86.6 | 57.1 | 92.5 | 92.7 |
|  | Outliers | 4.5 | 16.9 | 0 | 0 | 0 | 2.1 | 12.0 | 0.8 | 0.8 | 2.9 | 19.5 | 0.4 | 0.8 |
|  | Rama-Z | -4.5 | -8.1 | -3.5 | -2.7 | -1.3 | 0.8 | -7.3 | -3.3 | -2.3 | -4.4 | -7.4 | -3.5 | -2.9 |
| Rotamer outliers (%) |  | 5.5 | 19.2 | 5.5 | 6.9 | 4.1 | 18.1 | 26.4 | 7.3 | 7.3 | 8.7 | 27.4 | 2.1 | 3.3 |
| Clashscore |  | 50.8 | 43.9 | 3.5 | 1.4 | 1.4 | 14.3 | 69.8 | 6.3 | 1.9 | 17.8 | 89.1 | 2.4 | 2.8 |
| Cβ deviation | | 0 | 0 | 0 | 0 | 0 | 2.3 | 0 | 0 | 0 | 0 | 0 | 0 | 0 |

All refinements were performed using *qr.refine*. Quantum refinement was performed using the TeraChem (Ufimtsev & Martínez, 2009; Titov *et al.*, 2012) and xtb (Grimme *et al.*, 2017; Bannwarth *et al.*, 2019) packages as QM engines. Classic refinement (also referred here to as CCTBX refinement) used geometry restraints as implemented in CCTBX and used in *Phenix* refinement programs such as *phenix.refine* (Afonine *et al.*, 2012) and *phenix.real_space_refine* (Afonine, Poon, *et al.*, 2018). Classic refinement used only a standard set of geometry restraints: restraints on covalent bond lengths and bond angles, dihedral angles, chiralities, planarities and non-bonded repulsion (Engh & Huber, 1991, Grosse-Kunstleve *et al.*, 2002, Afonine *et al.*, 2004; Moriarty *et al.*, 2016). TeraChem refinement was performed using the Hartree-Fock (Slater, 1951) method with Grimme’s dispersion correction D3 (Grimme *et al.*, 2010) in conjunction with the 6-31G basis set (Hehre *et al.*, 1972) accounting for the polar (water, ε=78) environment by means of the COSMO polarizable continuum model (Liu *et al*, 2015, Barone & Cossi, 1998, Truong & Stefanovich, 1995). In what follows we will refer to this method as HF-D3/6-31G or simply TeraChem (TC). xtb refinement was performed with the original GFN-xTB model (Grimme *et al.*, 2017), nowadays denoted GFN1-xTB, and including the solvent effect by means of xtb's internal generalized Born solvation model. Atomic model fragmentation is the key step in the refinement procedure that makes QM calculations practical even for large molecules. Fragmentation as described in Zheng, Moriarty, *et al.* (2017) was used in all refinement runs. We refer to model geometry optimization (model regularization) as a special refinement regime using a modified version of (1) with the *T_data_* term entirely omitted.

Standard geometric criteria (Morris *et al.*, 1992) along with MolProbity (Williams *et al.*, 2018) tools were used to assess the quality of the atomic models before and after refinement. The distribution of the protein (φ, ψ) angles is indicative of refinement success and generally is sensitive to the model parameterization used (e.g., restraints). Thus, assessing Ramachandran plots is an integral part of model validation. However, comparison of two or more Ramachandran plots is not straightforward because dots on the plot that represent (φ, ψ) pairs are not labeled and for large models overlap often appearing as a single blob. Following Kleywegt & Jones (1996) we have implemented the *phenix.comparama* tool in *Phenix* that allows the tracking of changes of (φ, ψ) angles on the Ramachandran plot. The tool takes two models, for example, before and after refinement, and makes a Ramachandran plot for the latter model. Then for each (φ, ψ) pair that has moved from one region of the plot to another it draws an arrow that begins at the position of this pair in the starting model. Light green arrows show residues that left the outlier region, red arrows are for residues that became outliers, orange indicate favored-to-allowed transitions, and green stands for allowed-to-favored transitions.

## 3. Results and discussion

Table 1 summarizes data, model and model-to-data fit statistics for initial models and maps as obtained from PDB/EMDB along with the results of the refinement runs: quantum (TeraChem and/or xtb) and classic with two different ways to define the weight in (1). We observe that quantum refinement and optimization (see below, and also (Zheng *et al.*, 2020, Zheng, Reimers, *et al.*, 2017, Zheng, Moriarty, *et al.*, 2017) consistently produce models with RMSDs from library values for covalent bonds and angles of around 0.015 Å and 1.5 degrees, respectively, matching the community consensus on the targets for these metrics (see discussion and set of references in Afonine, P. V., Klaholz *et al*., 2018). At the same time low-resolution experimental data cannot justify these rather large deviations in these values, likely resulting in fitting the model to noise. This suggests that for low-resolution refinements the choice of weight in (1) should be such that the refinement produces models with bond and angle RMSDs as small as achievable in pure geometry regularization. With this in mind we performed two independent runs of classic refinement: one using the weight in (1) that results in bond and angle RMSDs of approximately 0.015 Å and 1.5 degrees (referred to as *CCTBX loose* in Table 1), and the other one that produces models with nearly idealized covalent geometry (referred to as *CCTBX tight* in Table 1).

Firstly, we note that all three initial models have rather poor geometry. Indeed, according to MolProbity (Chen *et al.*, 2010) a geometrically sound model is expected to have 98% or more residues in the favored region of the Ramachandran plot and less than 0.05% in disallowed, less than 0.3% of residue side chains being rotamer outliers, no C_β_ deviations and a low clashscore (the lower the better, typically 10-15 or below). None of the three models pass these criteria. The Ramachandran Z-score (Rama-Z) (Hooft *et al.*, 1997, van Beusekom *et al.*, 2018, Sobolev *et al.* 2020) provides a convenient numeric estimate of the protein main-chain conformation validity. The rule-of-thumb interpretation of the Ramachandran plot Z-score (Sobolev *et al.*, 2020) can be summarized as |Rama-Z| > 3 is poor indicating improbable backbone geometry, 2 < |Rama-Z| < 3 unlikely yet possible, and |Rama-Z| < 2 is good, normal backbone. Only one model, 5fn5, has this metric within the good range (0.8). However, considering the Rama-Z score separately for different secondary structure classes (such as helices, sheets and loops), we find that the Rama-Z score for helices is 1.6 (good) while it is −4.2 for loops (poor), with the standard deviations of 0.4 and 0.8, respectively. This means that overall the model main-chain conformation is problematic.

Figure 1 shows the Ramachandran plots for the deposited 3j63 as well as for re-refined models. As already stated, the deposited model (Fig. 1 a) is poor in both the Rama-Z score (−4.5) and percentage of residues in favored region of the plot (85.4). The TeraChem calculations transform these values to the “good” Rama-Z range (−1.3) and perfect percentage in favored region (Fig 1 b,c). The xtb model is also an improvement (−2.7, 98.9) on the initial model moving both statistics to the edge or into “good” regions in each case. The better quality of the TeraChem refinement comes with a multi-fold increased computational cost (from about 14 h for xtb on a 64-core CPU to 150 h for TeraChem on 8 nodes with 4 GPUs each).

**Figure 1.**
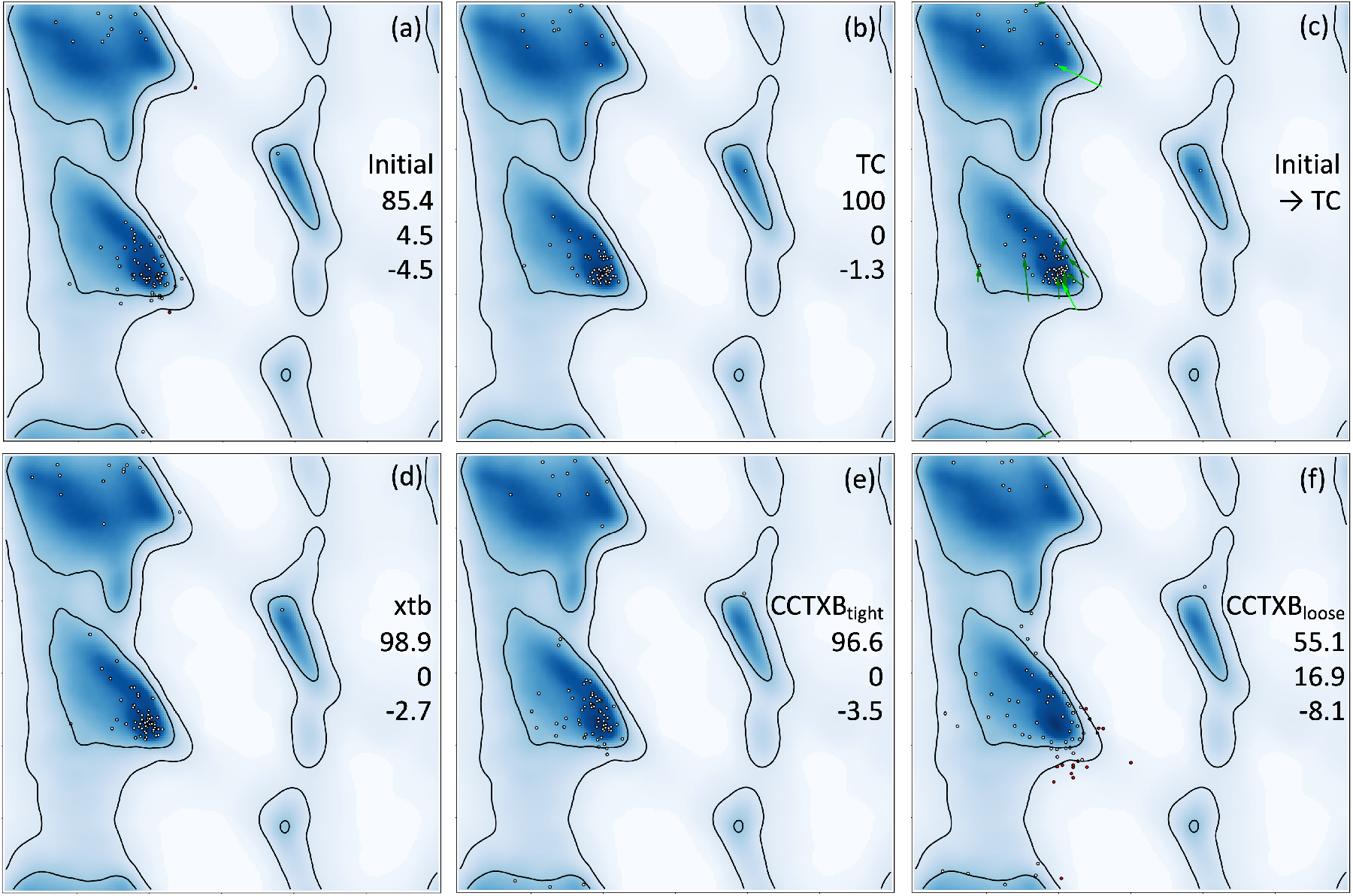
Ramachandran plots for 3j63: (a) initial; after quantum refinement using TeraChem (b,c) and xtb (d); and after classic refinement (CCTBX) using tight (e) and loose (f) geometry restraints. Columns of three numbers show percentage of residues in favored and disallowed regions of the plot, and the Rama-Z score. The Green and light-green arrows on the panel (c) show residues moved from allowed and disallowed regions to favored region after TC refinement.

Classic real-space refinement performs drastically different depending on the choice of the weight scale in (1). Relaxing geometry restraints (*CCTBX loose*) to produce comparable bond and angle deviations to quantum refinement results in models with very poor metrics such as Ramachandran plot (Fig. 1f), Rama-Z score, rotamers and clashscore (Table 1). Using very tight restraints (*CCTBX tight*) yields models with nearly idealized stereochemistry, while having unlikely secondary-structure geometry parameters as judged by the Ramachandran plot (Fig. 1e) and the Rama-Z score (Table 1).

Extending the xtb and classical refinements to include 3a5x and 5fn5 provides better statistics. The results covering the three models are shown in figure 2. The Rama-Z values for the xtb and classical refinements are similar to the single 3j63 model. Furthermore, for the xtb the distribution of residues on the Ramachandran plot is notably better as it follows the most probable regions in both alpha and beta segments of the plot (Fig. 2b). All the geometry parameter improvements occur at no expense to the model to data (map) fit, as indicated by the CC_mask_ values (Afonine *et al*., 2019) shown in table 1. Moreover, for 3a5x and 5fn5 quantum refinement significantly reduced the number of residue side-chain rotamer outliers compared to starting models. Fixing a rotamer outlier often requires traversing energy barriers in χ-angle space, which is unachievable by well-behaved gradient-driven optimization requiring additional methods such as simulated annealing (Brunger & Rice, 1997) or torsion exhaustive searches in χ-angle space (Oldfield, 2001). This explains the fact that none of the refinements produced outlier-free models. Also, we note that in some cases quantum refinement produced models with an elevated number of rotamer outliers compared to the *CCTBX tight* refinements. The reason for this is three-fold: 1) tight CCTBX restraints result in very strong nonbonded repulsions that force atoms into more favorable (but not necessarily most correct) conformations, 2) map quality in all three cases does not allow unambiguous placement of side-chains and as a result 3) allows hydrogen bond interactions between side-chain and neighbor atoms (often spurious due to poor initial model quality) that can generate sufficient strain forcing the side chain into a rotamer outlier.

**Figure 2.**
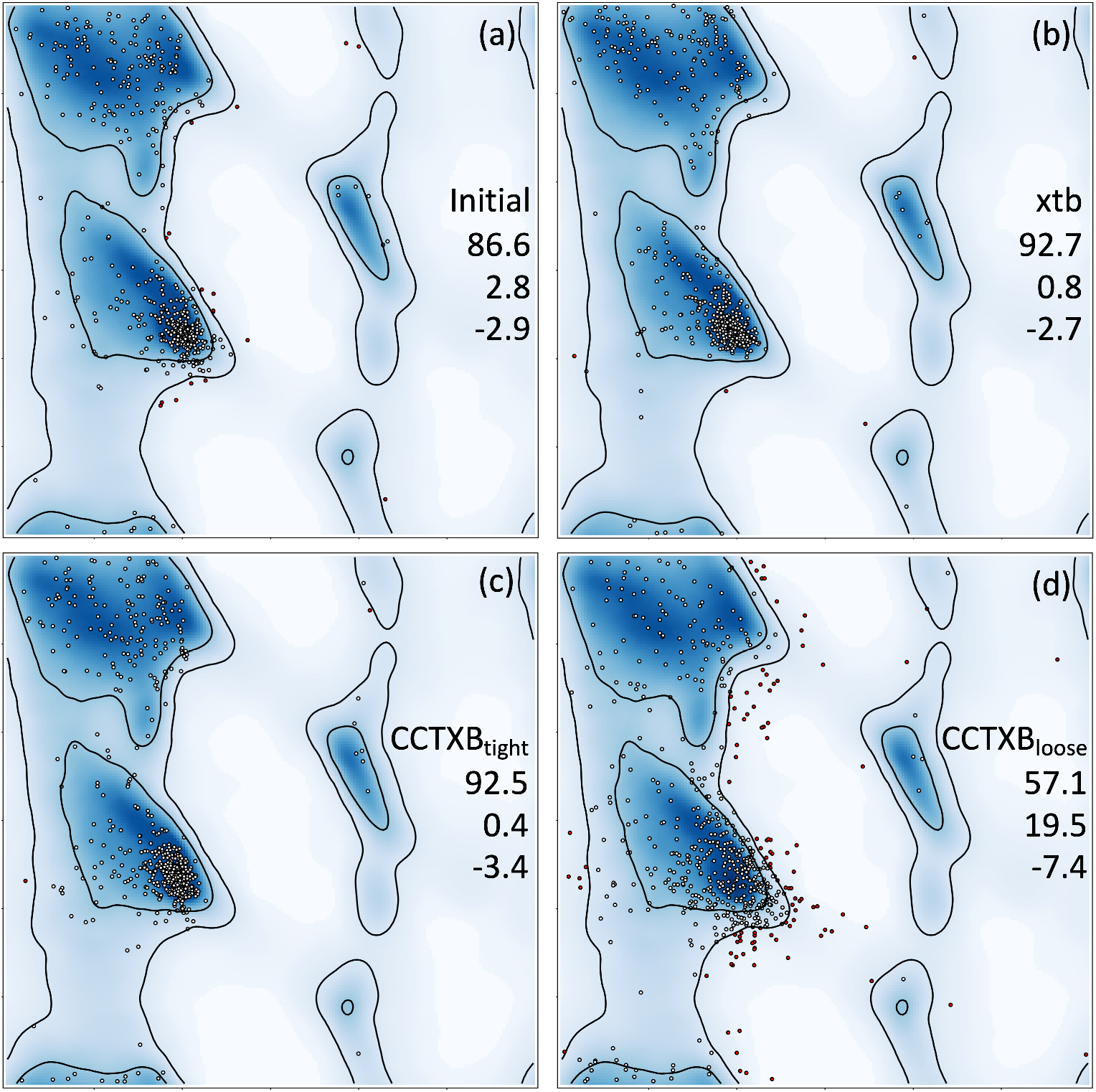
Each Ramachandran plot shows residues accumulated from all three models considered in this work (3j63, 5fn5 and 3a5x): (a) initial, (b) after quantum refinement (xtb) and after classic refinement (CCTBX) using tight (c) and loose (d) geometry restraints. Columns of three numbers show percentage of residues in favored and disallowed regions of the plot, and the Rama-Z score.

Expectedly, all classic refinements using loose restraints (*CCTBX loose*, Table 1) produce noticeably higher (better) map-model correlations CC_mask_ while also all of them show poorer model geometry. This can be interpreted as over-fitting of the data. We also note that classic refinements with tight restraints produce models with very similar CC_mask_ compared to xtb or TC. This means that these geometric improvements are on the scale of magnitude that can hardly be validated by the experimental data.

As follows from Table 1, the most notable improvement to the model resulting from quantum refinement was the quality of the protein main chain conformations. This is evidenced by significantly improved Ramachandran plot statistics, with significantly better Rama-Z scores obtained with TeraChem. The data resolution in all three selected examples is far from atomic and thus cannot validate this improvement. This led to the idea of performing an additional test that shows the superiority of QM restraints, and compares directly results from TeraChem and xtb. For this test we have selected one of the best-quality available model from the PDB solved at ultra-high resolution of 0.66 Å (1us0; Fig. 3a). For simplicity, we left out all non-protein atoms and atoms in alternative conformations (keeping just one conformer with the highest occupancy and resetting its occupancy to unity). Also, we created another copy of this model with perturbations of the atomic coordinates (Fig. 3b). We then subjected these two models to the pure geometry optimization using the TeraChem (HF-D3/6-31G), xtb (GFN1-xTB) packages and CCTBX, six optimizations were run in total. We rationalize this test as following. Given the high-resolution data used to determine the 1us0 model, we can assume that this model accurately represents the true structure. In turn, we expect that a better potential used to optimize the original model should not move or perturb it in any significant way, while the optimized perturbed model is expected to move back closer to the unperturbed original model. Indeed, Figure 3 shows that TeraChem based optimization brought the perturbed model very close to the original one (Fig. 3d) while not moving the original model significantly (Fig. 3c). The xtb approach yields some small deviations to the original model (Fig. 3e), but significantly improved the perturbed model (Fig. 3f). In contrast, optimization using classic restraints moved the perturbed model further away (Fig. 3h) from the original one. Moreover, the classic optimization substantially perturbed the original model (Fig. 3g). This test shows that replacement of classic restraints by the QM restraints result in more realistic molecular models even in the absence of experimental data.

**Figure 3.**
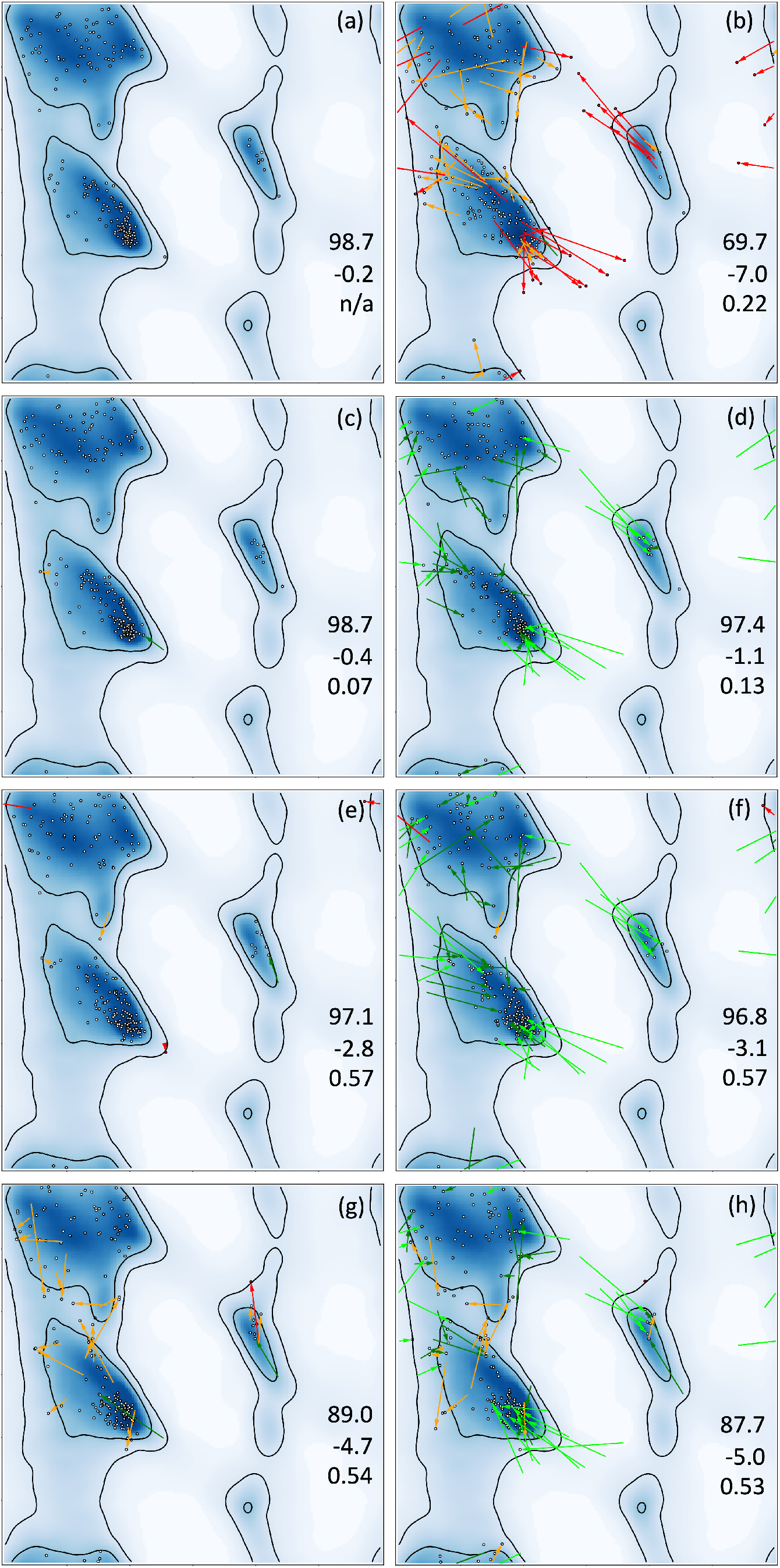
Ramachandran plots for 1us0: (a) original model, (b) perturbed model, (c, d) original and perturbed models, respectively, after QM optimization with TeraChem, (e, f) original and perturbed models, respectively, after QM optimization with xtb and (g, h) original and perturbed model, respectively, after classic optimization. Red and orange arrows show residues moved to disallowed and allowed regions out of favored region as result of perturbation. Green and light-green arrows show residues moved from allowed and disallowed regions to favored region. Columns of three numbers in each plot show percentage of residues in the favored region of the plot, Rama-Z score and RMSD (in Å) to the original (a) model. (b,c,d,e,f,g,h) plots were created using *phenix.comparama*.

## 4. Conclusions

Here we presented the new capability of the Q|R software package to perform real-space refinement enabling quantum refinement of cryo-EM derived models. The only input information required to run the refinement are the atomic model and the map. The atomic model is expected to be well-refined using standard classic refinement and be atom-complete. We once again demonstrated the superiority of the quantum-based restraints over the classic approach. In the three examples considered, we observe significant improvement of model geometry metrics after quantum refinement compared to the original models and to models after classic refinements. The HF-D3/6-31G level of theory performs better than the GFN1-xTB model for our refinements. A new tool for graphical comparison of Ramachandran plots (*phenix.comparama*) has been developed to aid this work and is now part of *Phenix* software suite.

Using quantum refinement from Q|R project requires installing Q|R software (www.qrefine.com), *Phenix* (www.phenix-online.org) and one of the supported quantum refinement programs, such as TeraChem, Orca, or xtb.

Re-refinement results and related data are available at: https://github.com/qrefine/QR3-RSR-cryo-EM.

## Acknowledgments

This work was enabled by the financial support from the National Natural Science Foundation of China (Grant No. 31870738) and support from the Shanghai Eastern Scholar Program. HK acknowledges support by the ERDF project SYMBIT (CZ.02.1.01/0.0/0.0/15_003/0000477).

## Footnotes

1 Different sources quote different threshold for defining ultra-high resolution on seemingly arbitrary grounds. Here we use 0.7 Å as described by Urzhumtsev *et al.* (2009) because they seem to provide most solid rationale for the choice of this resolution cutoff.

